# Network pharmacological evaluation of active ingredients and potential targets of San Huang Decoction in the treatment of breast cancer

**DOI:** 10.1101/2024.06.20.599797

**Authors:** Shuai Chao, Shiqiang Liu, Kun Wang, Mingming Xie, Beibei Liu

## Abstract

**Objective:** Using the method of network pharmacology, the possible targets and related signaling pathways of San Huang Decoction against breast cancer were explored.

**Methods:** The active ingredients of San Huang Decoction were screened in Traditional Chinese Medicine Systems Pharmacology Database and Analysis Platform (TCMSP). All targets of San Huang Decoction and breast cancer were obtained through five databases, and R software was used to obtain their common targets. Then, Cytoscape software was used to construct the active ingredient-target network map. The protein-protein interaction network (PPI) was obtained with the help of the STRING database, and key targets were screened out through network topology analysis. Common targets were subjected to gene ontology (GO) and Kyoto encyclopedia of genes and genomes (KEGG) enrichment analysis using R software.

**Results:** A total of 39 active ingredients were screened from San Huang Decoction, of which 20 belonged to Huangqi, 16 belonged to Dahuang, and 3 belonged to Jianghuang. After taking the intersection of all the targets of San Huang Decoction and breast cancer, a total of 185 potential targets of San Huang Decoction against breast cancer were obtained. PPI network analysis screened out 16 targets that may be the key targets of San Huang Decoction against breast cancer. The enrichment analysis of GO and KEGG suggested that the common targets were involved in various biological processes and functions through PI3K-Akt signaling pathway, MAPK signaling pathway, TNF signaling pathway, IL-17 signaling pathway, and HIF-1 signaling pathway, including oxidation stress, transcription-related functions, and nuclear receptor functions.

**Conclusion:** This study predicted the potential targets and signaling pathways of San Huang Decoction against breast cancer, and provided scientific guidance for the further study on the anti-breast cancer mechanism of San Huang Decoction.

## 1 Introduction

According to the latest global cancer data released by the International Agency for Research on Cancer in 2020, breast cancer has replaced lung cancer as the world’s most morbidity cancer, and its mortality rate ranks first among female cancer deaths. In China, the incidence of breast cancer is characterized by the early age of onset, the relatively late clinical stage at the time of diagnosis, and the lower survival period than in European and American countries (1). Currently, breast cancer therapies include surgery, radiotherapy and chemotherapy, targeted therapy, and immunotherapy (2). However, for patients with recurrence and metastasis, the development of treatment methods cannot solve the problem of high breast cancer mortality (3). Therefore, more effective treatments need to be sought.

San Huang Decoction is a traditional Chinese medicine compound consisting of Huangqi, Dahuang, and Jianghuang (4). Previous studies have shown that these three Chinese herbal medicines have a regulatory effect on cancer growth. Xu et al. (5) found that Huangqi could promote the apoptosis of lung cancer cells. Zhang et al. (6) revealed that Dahuang had inhibitory effectcomm on hepatoma cells proliferation. Aggarwal and his colleagues had shown the anti-inflammatory and anti-cancer effects of Jianghuang (7). In addition, recent studies have shown that San Huang Decoction has an intervening effect on breast cancer growth and inflammation (8, 9). Although some progress has been made in the pharmacological research of San Huang Decoction and its monomers, the research on its molecular biology is relatively scarce.

Network pharmacology is based on the similarities between drugs and drugs in terms of structure and efficacy, and considers the multiple interactions of target molecules and biological effect molecules in the body, and conducts joint analysis of disease-related genes to construct Drug-component-target-disease regulatory network (10, 11). Chinese medicine network pharmacology emphasizes the multi-channel regulation of signal pathways, improves the therapeutic effect of drugs, reduces toxic and side effects, thereby increasing the success rate of clinical trials of new drugs and saving drug R&D costs (12). This study aims to probe the potential active ingredients, target genes and mechanisms of San Huang Decoction in breast cancer treatment through network pharmacology.

## 2 Materials and methods

### 2.1 Composition and screening the active ingredients of San Huang Decoction

The composition of San Huang Decoction is as follows: Huangqi, 30 g; Dahuang, 10 g; Jianghuang, 10 g (4). Traditional Chinese Medicine Systems Pharmacology Database and Analysis Platform (TCMSP, http://tcmspw.com/tcmsp.php) database was used to collect the ingredients and targets of the three herbs in San Huang Decoction. The screening conditions of active ingredients in the database were oral bioavailability (OB) ≥ 30% and drug-likeness (DL) ≥ 0.18.

### 2.2 Construction of disease related genes

Breast cancer-related genes were collected from GeneCards (https://www.genecards.org/), OMIM (Online Mendelian Inheritance in Man, https://omim.org/), PharmGkb (https://www.pharmgkb.org/), TTD (Therapeutic Target Database, http://db.idrblab.net/ttd/) and DrugBank (https://www.drugbank.ca/) databases. We searched the above database with the keyword “breast cancer” and downloaded the data. Then, the downloaded data were integrated using R language to draw a Venn diagram and get a file of disease-related genes. The venn package was utilized.

### 2.3 Intersection of targets of San Huang Decoction and breast cancer-related genes

R language was used to analyze the files of targets of San Huang Decoction and breast cancer-related genes, and finally generate a Venn diagram and a file of common target genes. The venn package was utilized.

### 2.4 Construction of compound-target network

Cytoscape software (version 3.8.2) was used to construct a visual compound-target network to intuitively reflect the relationship between active ingredients of San Huang Decoction and common target genes (mentioned in section 2.3).

### 2.5 Protein–protein interaction (PPI) network and topological analysis

The common target genes obtained above were uploaded to STRING online software (https://string-db.org/) to build a PPI network and download a TSV file. The PPI network filtering conditions include minimum required interaction score greater than 0.9 and hidden disconnected nodes in the network. The cytoNAC plugin was installed in Cytoscape software (version 3.8.2), and then the above TSV file was imported in the software for topology analysis. The analysis mainly focuses on the parameters of betweenness centrality (BC), closeness centrality (CC), degree centrality (DC), eigenvector centrality (EC), local average connectivity-based method (LAC), and network centrality (NC). Finally, the relevant network diagrams and files were exported.

### 2.6 Enrichment analysis of gene ontology (GO) and Kyoto encyclopedia of genes and genomes (KEGG)

The “colorspace,” “stringi,” and “ggplot2” packages were installed in R software. GO and KEGG enrichment analysis was performed by BiocManager package includes “clusterProfiler,” “DOSE,” “enrichplot”, and “pathview”. The filtration conditions of p value and q value were both 0.5. GO is divided into three parts: molecular function (MF), biological process (BP) and cellular component (CC). According to the count score, the top 10 of each part were selected for presentation (bar plot and bubble plot). For KEGG enrichment analysis, the top 30 were selected for display (bar plot and bubble plot) according to the count score. The package “pathview” was used to draw KEGG pathway map.

## 3 Results

### 3.1. Active ingredients of San Huang Decoction

According to the TCMSP database, San Huang Decoction contains a total of 231 ingredients, including 87 in Huangqi, 92 in Dahuang, and 52 in Jianghuang. Through the screening conditions of OB ≥ 30% and DL ≥ 0.18, 39 active ingredients were identified: 20 in Huangqi, 16 in Dahuang, and 3 in Jianghuang (Table 1).

**Table 1:**
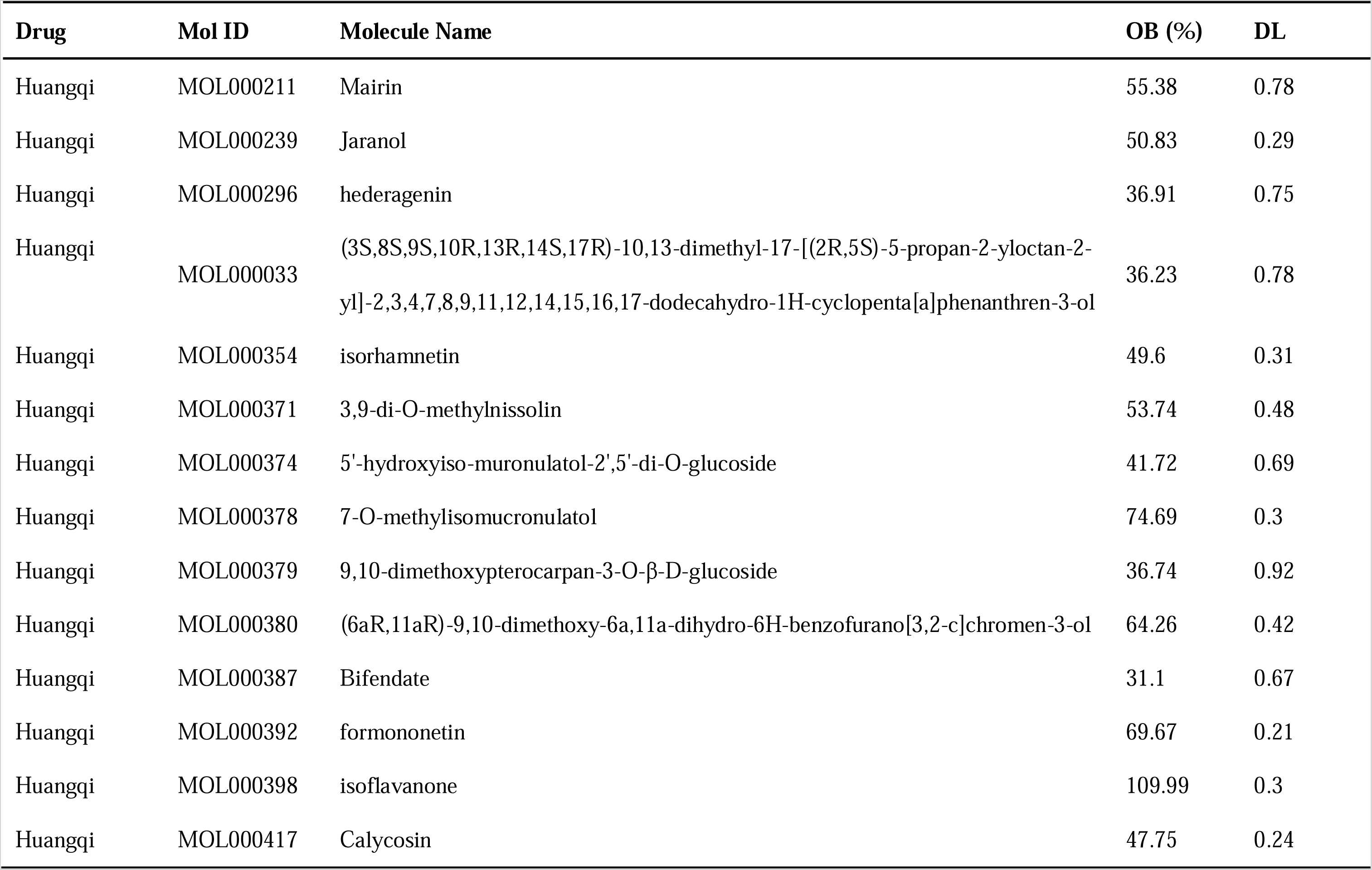
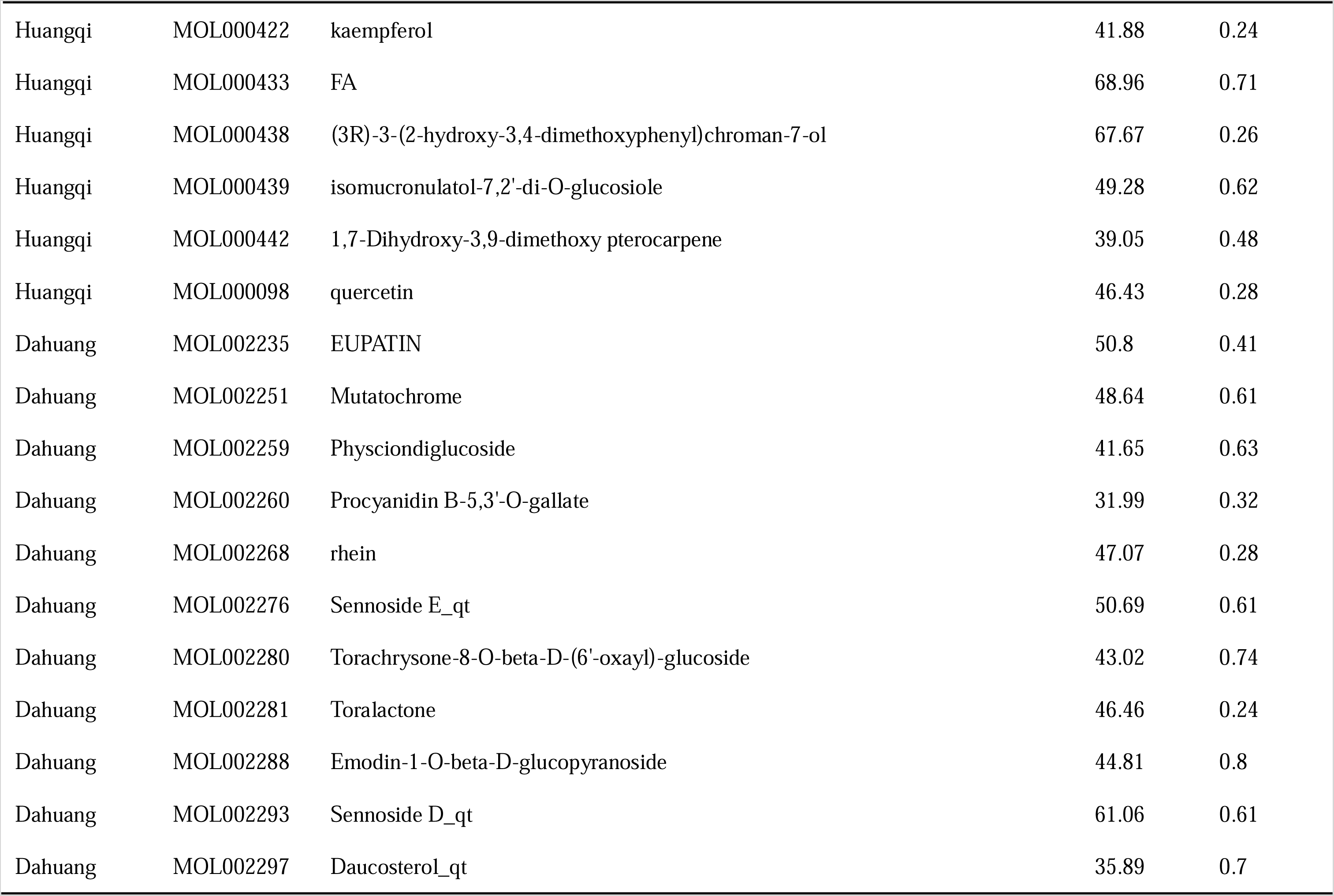
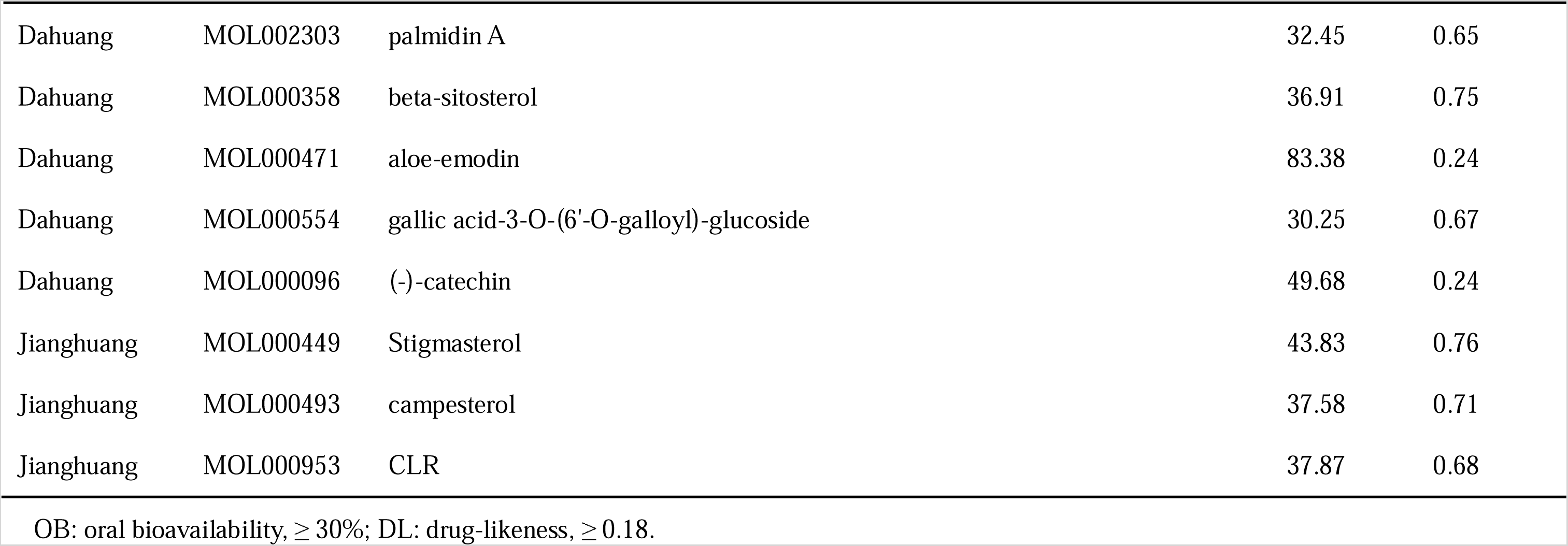
The active ingredients of San Huang Decoction.

### 3.2 The prediction and analysis of targets

Through the method of multi-source database integration, the related gene data of breast cancer from GeneCards, OMIM, PharmGkb, TTD and DrugBank databases were integrated. A total of 11673 breast cancer related genes were obtained, including 10938 genes from GeneCards, 168 from OMIM, 289 from PharmGkb, 135 from TTD, and 143 from DrugBank (Figure 1A). Then, through the intersection of drug targets and disease-related genes, 185 common targets were finally obtained (Figure 1B) for the further mechanism research of San Huang Decoction in the treatment of breast cancer.

**Figure 1:**
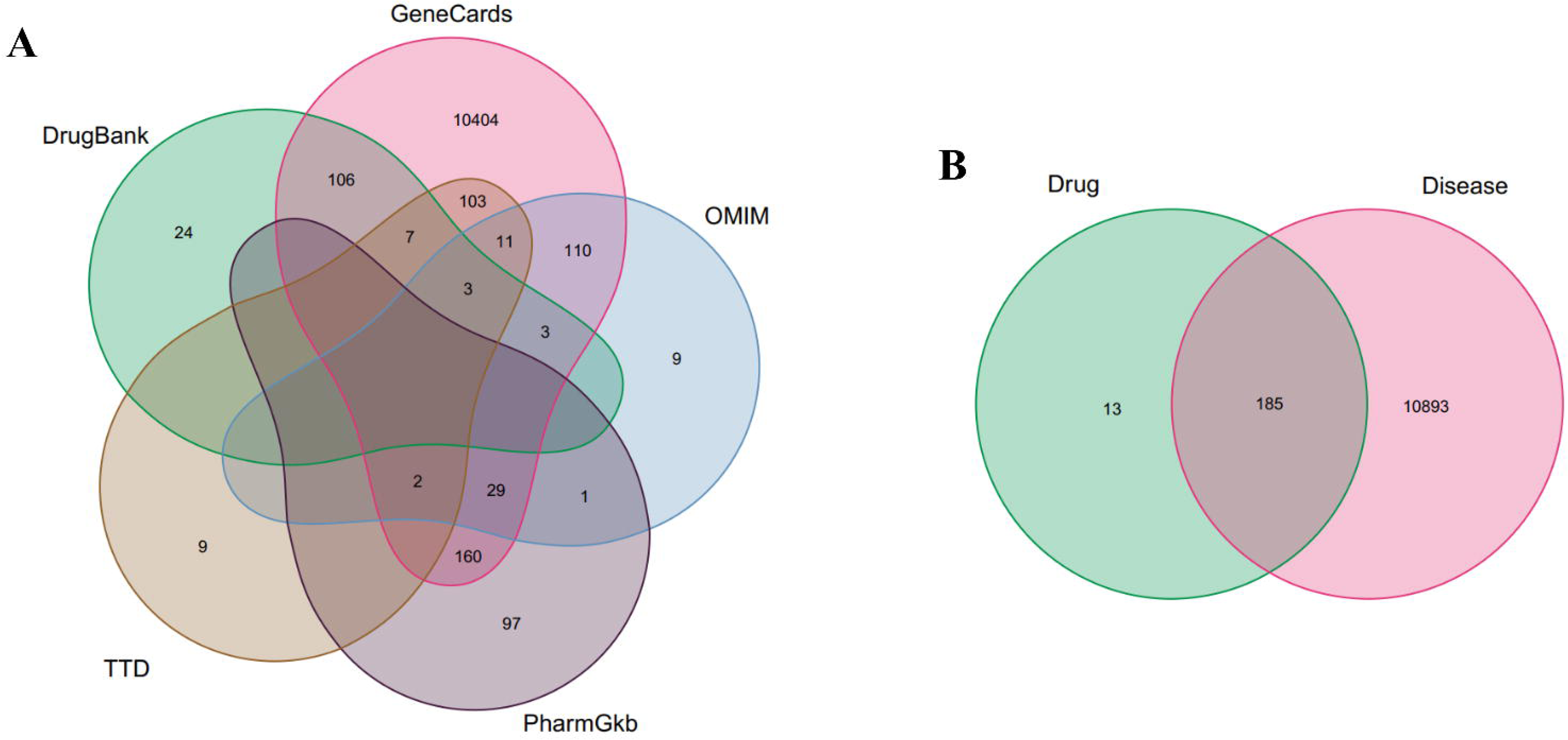
Venn diagram of the target. (A) The Venn diagram of GeneCards, OMIM, PharmGkb, TTD and DrugBank databases. (B) The Venn diagram of San Huang Decoction target and breast cancer target.

### 3.3 Compound-target network constructed and PPI network analysis

To clarify the relationship between target genes and compounds, we deleted the non-target compounds and constructed the compound-target network of San Huang Decoction by using Cytoscape software. The network consisted of 211 nodes and 473 edges (Figure 2). Then, 185 common targets were analyzed through STRING online software, and a PPI network with a minimum required interaction score greater than 0.9 was constructed (disconnected nodes in the network are hidden). The network included 185 nodes and 667 edges (Figure 3). The average node degree was 7.21. Subsequently, the topology analysis of the PPI network was performed. The median value of the three main parameters of “BC”, “DC” and “CC” was taken to construct the central node of the anticancer effect of San Huang Decoction on breast cancer. The filter value of the first screening was BC ≥ 90.38891286, DC ≥ 6 and CC ≥ 0.112982512, and 46 nodes and 257 edges were obtained (Figure 4). Then, the 46 nodes were screened for the second time. The filter value was BC ≥ 23.48603965, DC ≥ 10 and CC ≥ 0.5325630255, and finally 16 central nodes and 72 edges were obtained (Figure 4). The 16 central nodes included TP53, RB1, MAPK14, CCND1, MYC, MAPK1, AKT1, CDK1, ESR1, CDKN1A, JUN, FOS, EGFR, NFKBIA, MAPK8, and RELA (Table 2).

**Figure 2:**
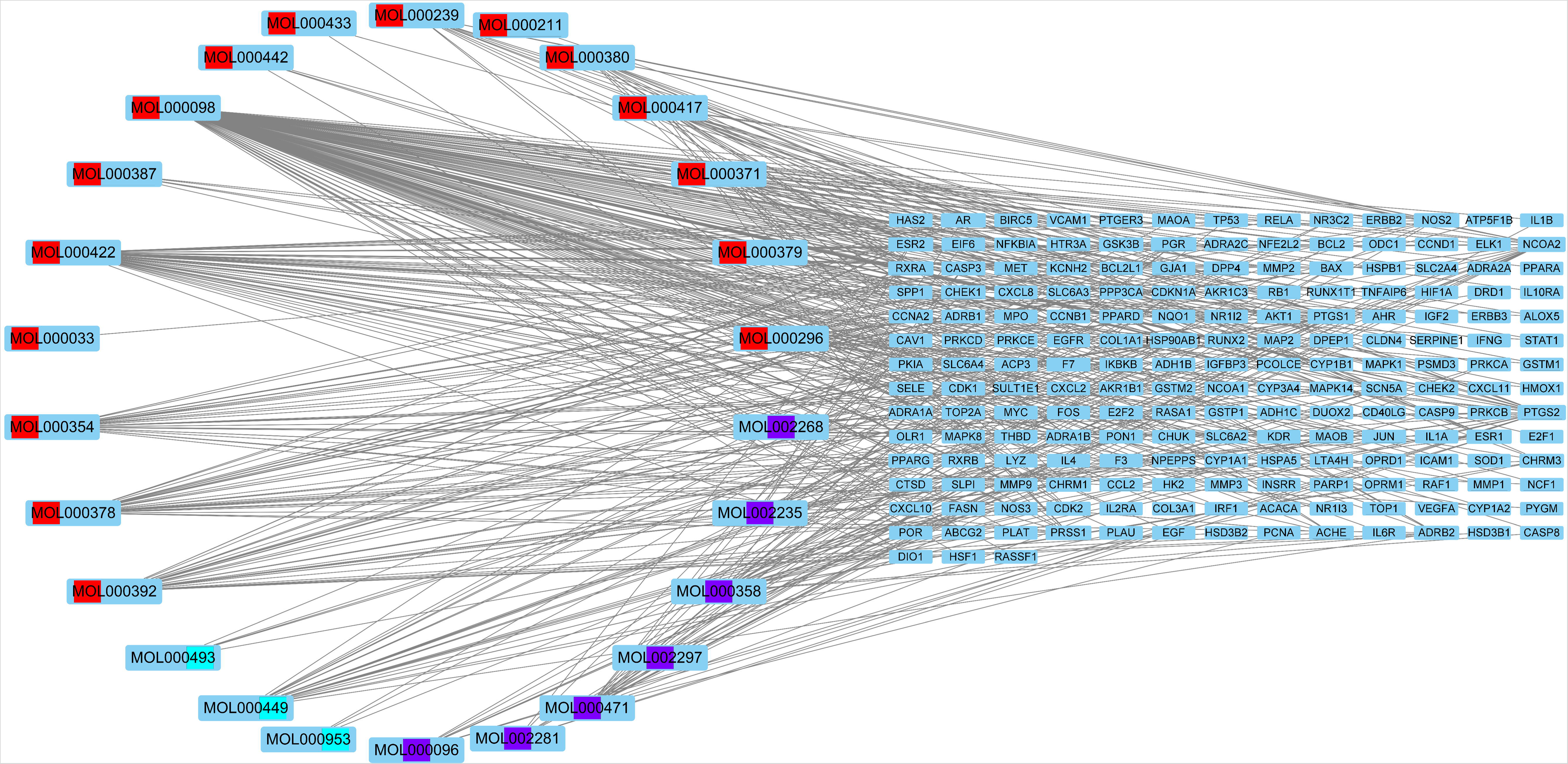
Compound target network of San Huang Decoction. Red: Huangqi, Purple: Dahuang, Baby blue: Jianghuang.

**Figure 3:**
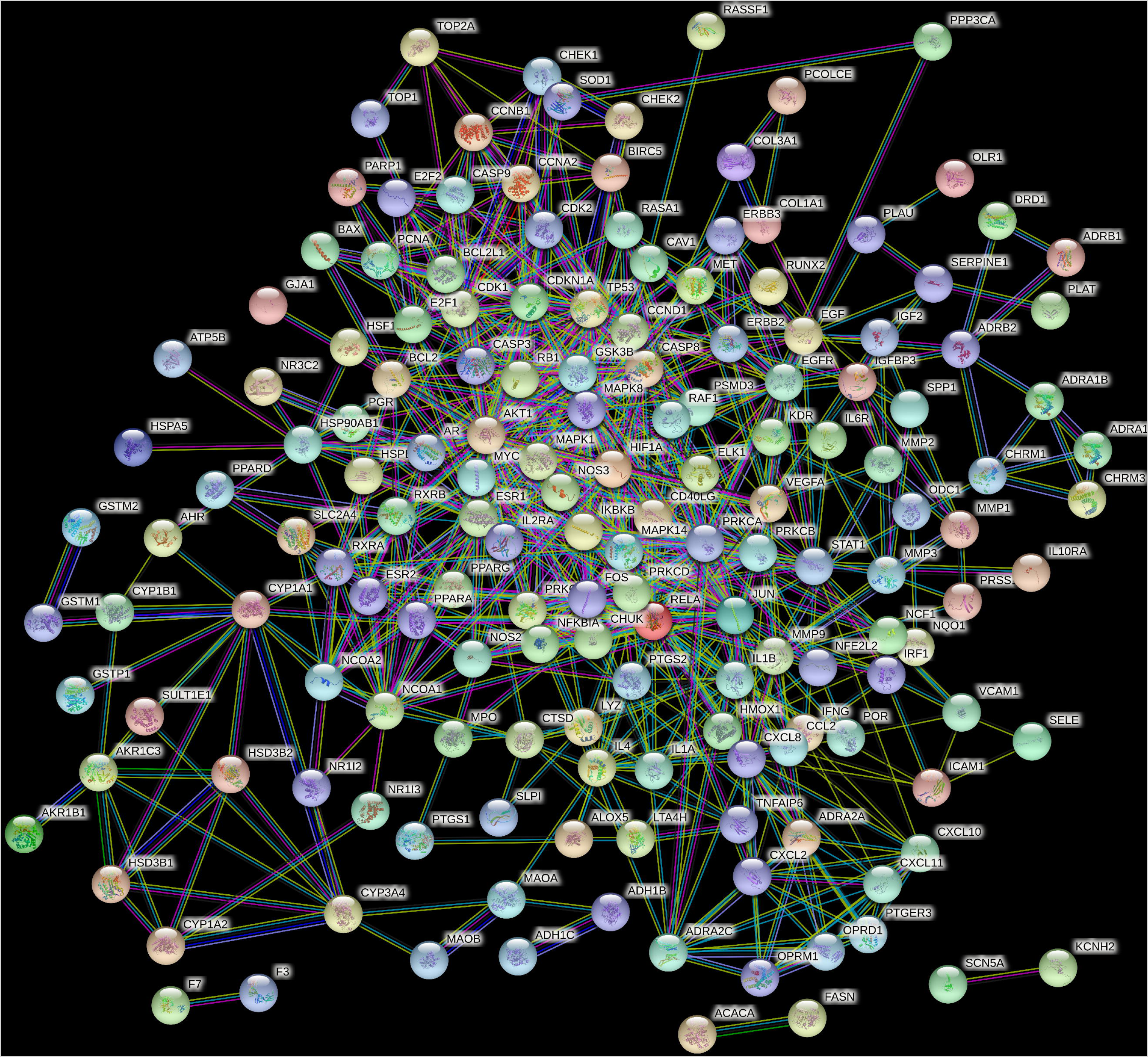
PPI network analysis of target proteins. Network nodes represent proteins. Edges represent protein-protein associations. The protein structures are displayed in the nodes.

**Figure 4:**
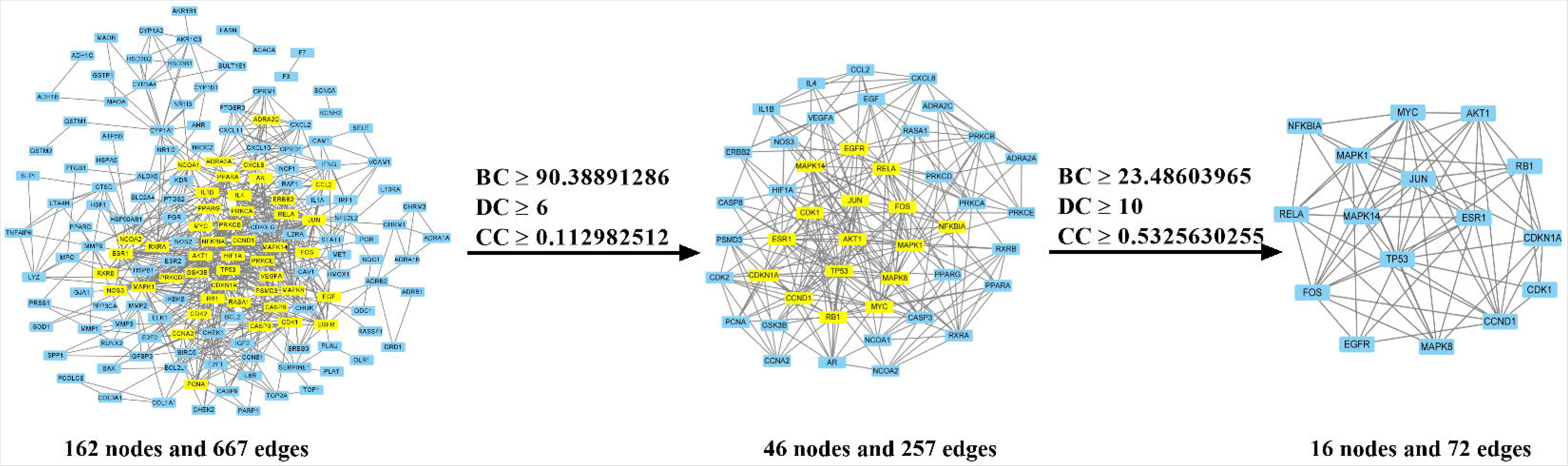
The process of topological screening for PPI network.

**Table 2:** Results of topology analysis.

| **Gene symbol** | **Description** | **BC** | **DC** | **CC** |
| --- | --- | --- | --- | --- |
| TP53 | Tumor protein P53 | 212.6503132 | 26 | 0.703125 |
| RB1 | RB transcriptional corepressor 1 | 50.7647377 | 15 | 0.584415584 |
| MAPK14 | Mitogen-activated protein kinase 14 | 59.39869723 | 15 | 0.592105263 |
| CCND1 | Cyclin D1 | 43.99401849 | 16 | 0.592105263 |
| MYC | MYC proto-oncogene | 29.81463281 | 15 | 0.584415584 |
| MAPK1 | Mitogen-activated protein kinase 1 | 164.5715737 | 22 | 0.661764706 |
| AKT1 | AKT serine/threonine kinase 1 | 96.34125785 | 18 | 0.608108108 |
| CDK1 | Cyclin dependent kinase 1 | 47.69657504 | 11 | 0.535714286 |
| ESR1 | Estrogen receptor 1 | 61.91770757 | 15 | 0.584415584 |
| CDKN1A | Cyclin dependent kinase inhibitor 1A | 28.8292103 | 14 | 0.5625 |
| JUN | Jun proto-oncogene | 188.5643286 | 24 | 0.681818182 |
| FOS | Fos proto-oncogene | 26.17196219 | 13 | 0.569620253 |
| EGFR | Epidermal growth factor receptor | 40.37580475 | 12 | 0.5625 |
| NFKBIA | NFKB inhibitor alpha | 24.28786982 | 11 | 0.555555556 |
| MAPK8 | Mitogen-activated protein kinase 8 | 56.25016687 | 14 | 0.592105263 |
| RELA | RELA proto-oncogene | 137.8152165 | 19 | 0.633802817 |

BC: betweenness centrality; DC: degree centrality; CC: closeness centrality.

### 3.4 Enrichment analysis of GO and KEGG

To understand the mechanism of the active ingredients of San Huang Decoction on breast cancer, GO enrichment analysis was performed on 185 common targets in Figure 1B. GO is divided into three parts: MF, BP and CC. Both the bar plot (Figure 5A) and bubble plot (Figure 5B) showed the top ten terms of each part. To reveal the main pathways of the 185 common targets, we performed KEGG enrichment analysis. As a result, the top 30 terms were screened out and a bar plot (Figure 6A) and bubble plot (Figure 6B) were drawn. These pathways included AGE-RAGE signaling pathway in diabetic complications (hsa04933), fluid shear stress and atherosclerosis (hsa05418), IL-17 signaling pathway (hsa04657), kaposi sarcoma-associated herpesvirus infection (hsa05167), human cytomegalovirus infection (hsa05163), TNF signaling pathway (hsa04668), endocrine resistance (hsa01522), Th17 cell differentiation (hsa04659), Epstein-Barr virus infection (hsa05169), proteoglycans in cancer (hsa05205), cellular senescence (hsa04218), PI3K-Akt signaling pathway (hsa04151), EGFR tyrosine kinase inhibitor resistance (hsa01521), human T-cell leukemia virus 1 infection (hsa05166), HIF-1 signaling pathway (hsa04066), platinum drug resistance (hsa01524), p53 signaling pathway (hsa04115), and MAPK signaling pathway (hsa04010). Moreover, prostate cancer (hsa05215), hepatitis B (hsa05161), bladder cancer (hsa05219), hepatitis C (hsa05160), pancreatic cancer (hsa05212), small cell lung cancer (hsa05222), non-small cell lung cancer (hsa05223), hepatocellular carcinoma (hsa05225), measles (hsa05162), colorectal cancer (hsa05210), toxoplasmosis (hsa05145), and breast cancer (hsa05224) are also highly correlated with key target genes. Figure 7 showed a diagram of the KEGG pathway in breast cancer. White labels indicate other related targets or signal pathways. Red labels indicate the target or signal pathway related to San Huang Decoction.

**Figure 5:**
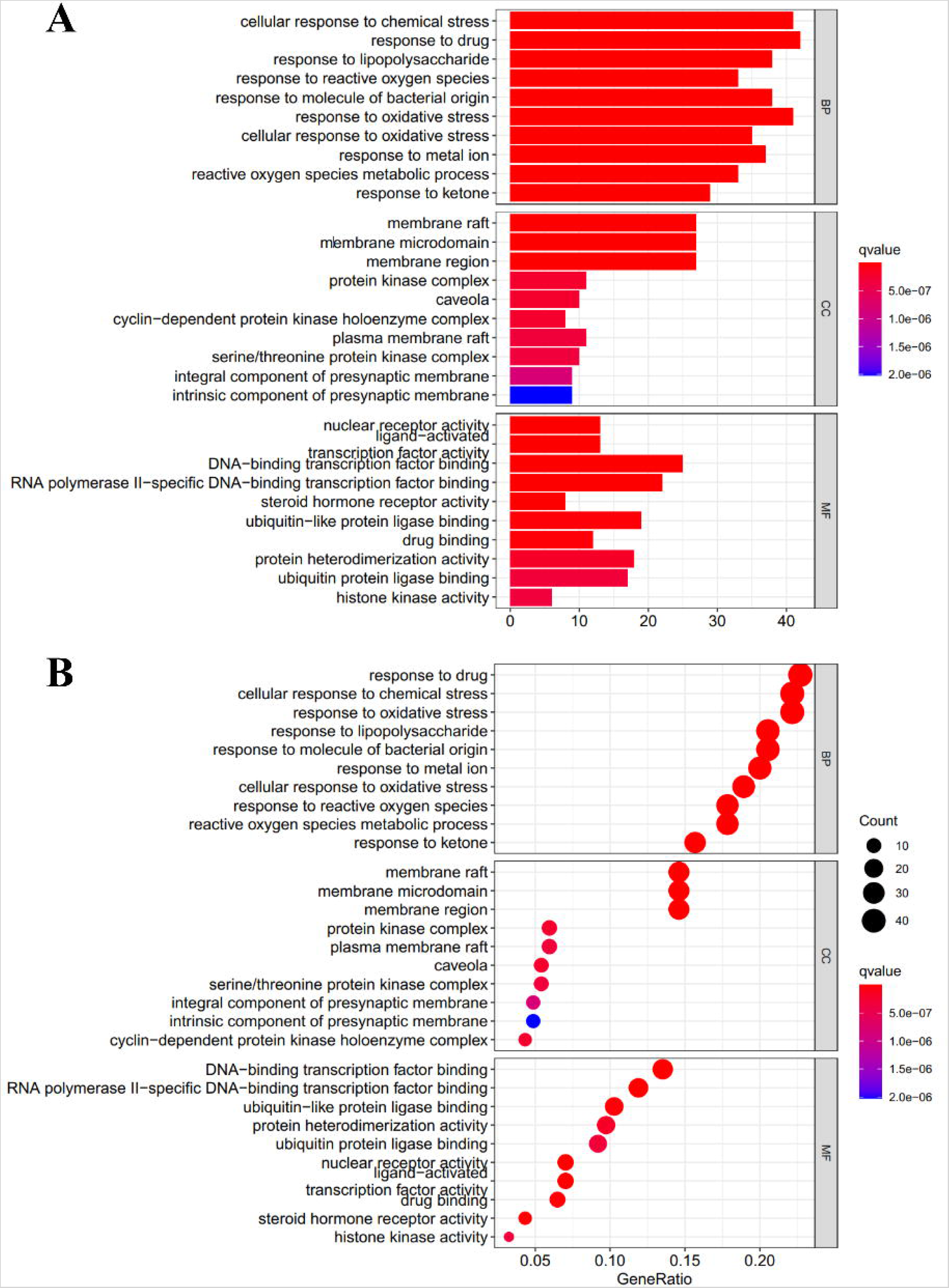
GO enrichment analysis. (A) The bar plot of GO enrichment analysis. (B) The bubble plot of GO enrichment analysis. The top 10 terms of BP, MM, and CC were selected to demonstrate. The color of terms turned from blue to red. In the bubble plot, the larger the size of the dot, the more the number of enriched genes, and the more the color of the dot is red, the more significant the enrichment.

**Figure 6:**
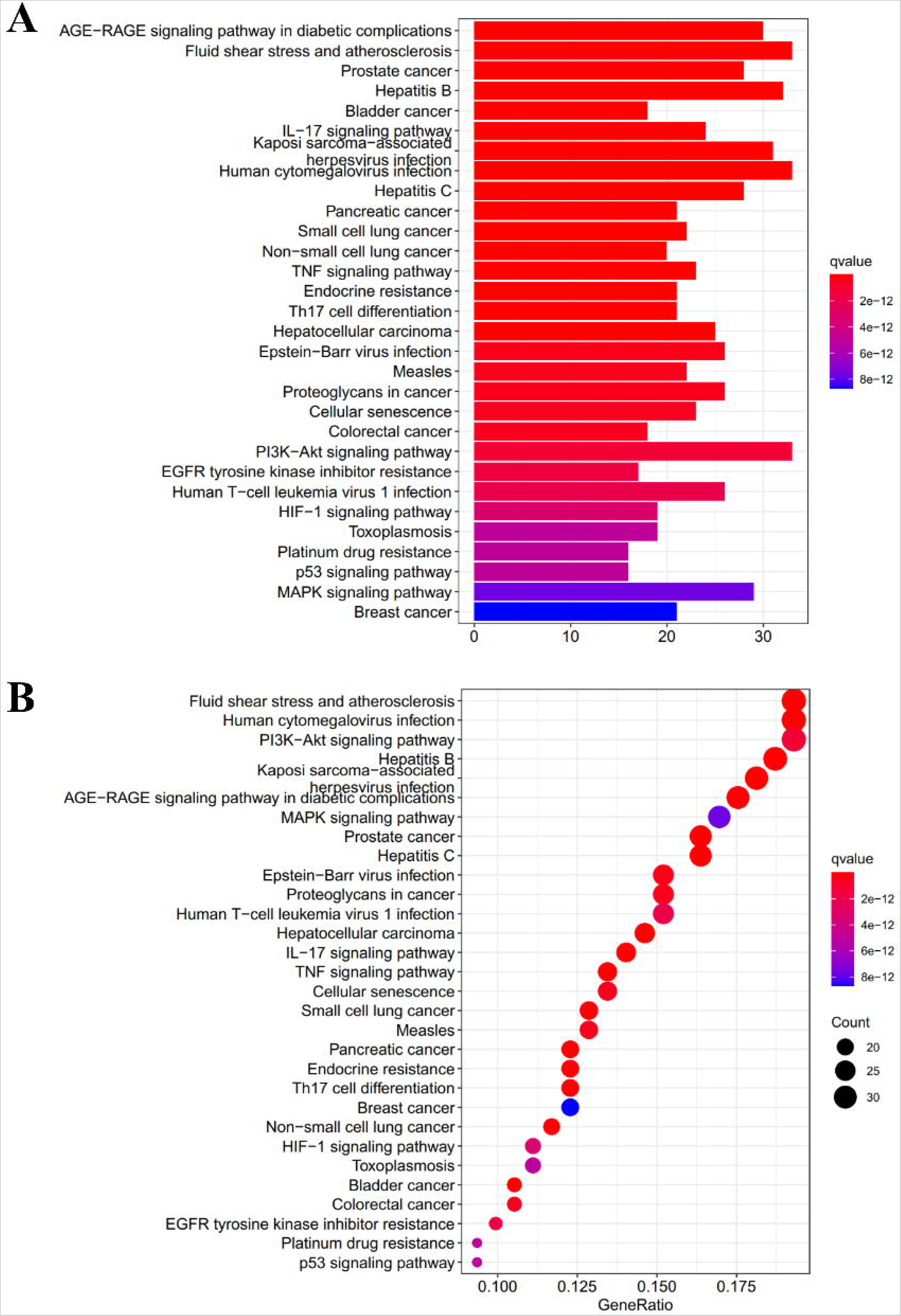
KEGG enrichment analysis. (A) The bar plot of KEGG enrichment analysis. (B) The bubble plot of KEGG enrichment analysis. The top 30 terms were selected to demonstrate. The color of terms turned from blue to red. In the bubble plot, the larger the size of the dot, the more the number of enriched genes, and the more the color of the dot is red, the more significant the enrichment.

**Figure 7:**
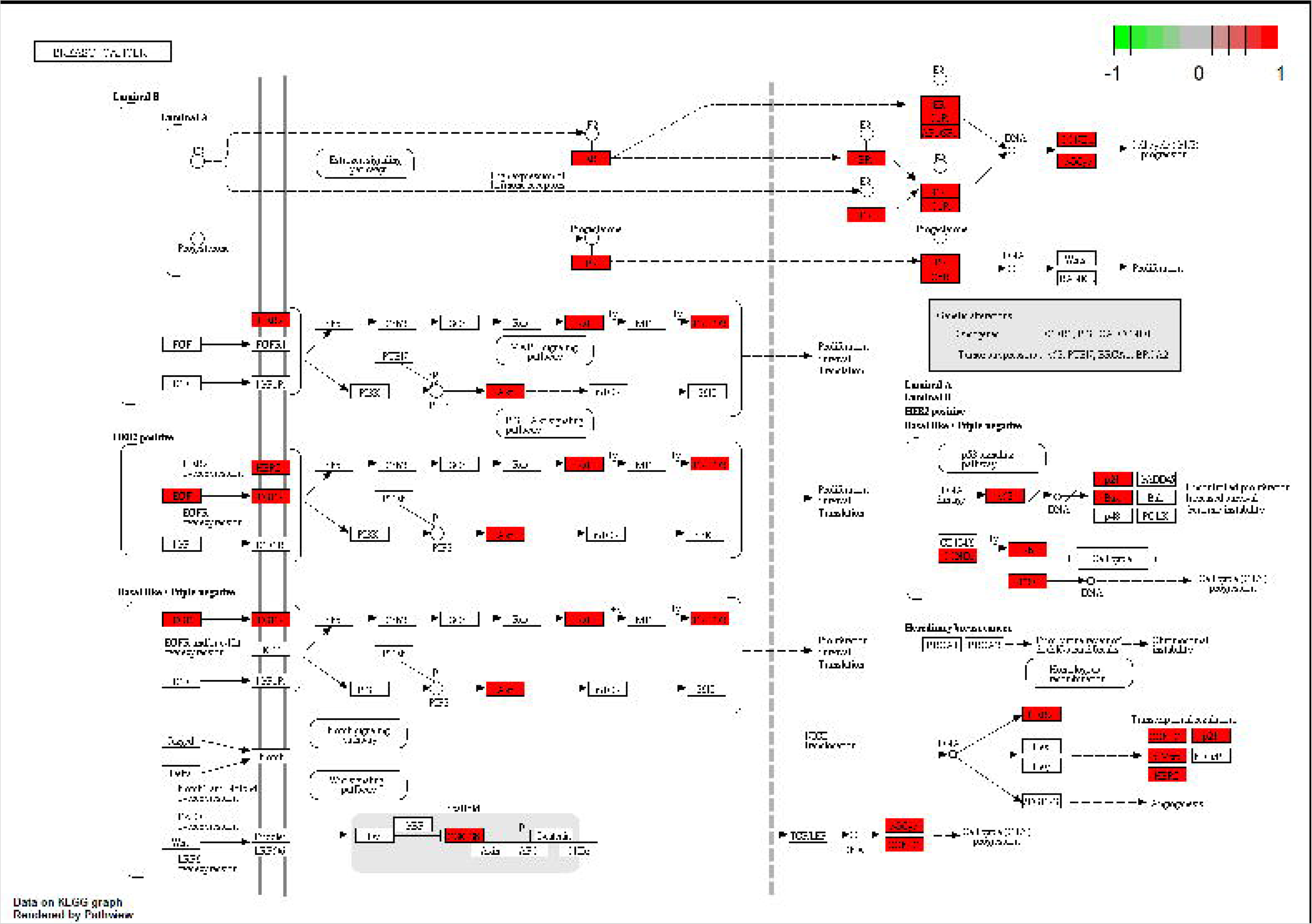
KEGG pathway in breast cancer.

## 4 Discussion

Despite numerous treatments for breast cancer, it remains the leading cause of cancer death in women worldwide. Therefore, finding new drugs to treat breast cancer is of great significance for clinical treatment. In recent years, San Huang Decoction has been used in the clinical treatment of breast cancer patients (8). Although some studies have shown that San Huang Decoction can inhibit the growth of breast cancer cells (4), the underlying mechanism of San Huang Decoction in the treatment of breast cancer is still unclear. In this study, we employed a network pharmacology-based approach to reveal the pharmacological mechanism of San Huang decoction against breast cancer, which may provide new therapeutic strategies for better treatment of breast cancer.

We obtained the chemical components in San Huang Decoction through the TCMSP database, and screened out 39 active ingredients according to the filter conditions of OB≥30% and DL≥0.18, such as Hederagenin, Bifendate, Mairin, isorhamnetin, Jaranol, isoflavanone, kaempferol, formononetin, Calycosin, 3,9-di-O-methylnissolin, quercetin, Mutatochrome, Physciondiglucoside, rhein, Toralactone, and Stigmasterol. These active ingredients are associated with anticancer effect. For instance, Kaempferol has been shown to have multiple biological effects, including inhibiting cell proliferation of various cancers, and participating in anticancer signal transduction pathways (13). Hederagenin enhanced the cytotoxicity of cisplatin and paclitaxel on lung cancer cells by blocking autophagy (14). Isorhamnetin inhibited the proliferation and metastasis of human gallbladder cancer cells through PI3K/AKT signaling pathway (15). Moreover, several studies have shown that Formononetin can inhibit the growth of breast cancer cells (16–18).

Next, we collected breast cancer-related target genes by merging data from five databases, and obtained 185 common genes by intersecting these target genes and the targets of San Huang Decoction. PPI network was constructed for these genes with STRING, and topology analysis was performed. Finally, 16 core genes of San Huang Decoction against breast cancer were screened from these 185 common genes, including TP53, RB1, MAPK14, CCND1, MYC, MAPK1, AKT1, CDK1, ESR1, CDKN1A, JUN, FOS, EGFR, NFKBIA, MAPK8, and RELA. Among them, TP53 is a transcription factor that can inhibit cell division or survival. Its mutations have been associated with a variety of human cancers (19, 20), including breast cancer (21). MYC functions primarily as a transcriptional regulator that regulates many different cellular processes, including cell growth, cell differentiation, angiogenesis, and immune responses (22). One study reports that USP29t coordinates the stabilization of MYC and HIF1α to promote cancer cell growth and metabolism (23). ESR1 is an estrogen receptor that combines with estrogen *in vivo*, promotes breast cell proliferation and differentiation, and is also involved in the pathological process of breast cancer (24) or endometrial cancer (25).

Go and KEGG enrichment analysis showed that these 185 common genes were mainly related to some signaling pathways, such as PI3K-Akt signaling pathway, MAPK signaling pathway, TNF signaling pathway, IL-17 signaling pathway, and HIF-1 signaling pathway. As one of the important signal transduction pathways in cells, PI3K-Akt signaling pathway plays a key role in inhibiting apoptosis and promoting proliferation in cells by affecting the activation state of various downstream effector molecules.It is closely related to the occurrence and development of various human tumors (26). The active ingredient isorhamnetin in San Huang Decoction inhibits the proliferation and metastasis of human gallbladder cancer cells through PI3K-Akt signaling pathway (15). Kaempferol is an active ingredient that promotes apoptosis of human cervical cancer cells through PI3K-Akt signaling pathway (27). MAPK signaling pathway is one of the important signal transduction systems in living organisms, involved in mediating various physiological and pathological processes such as cell growth, development and differentiation. MAPK cascade is a key pathway for cancer cell survival, propagation and drug resistance treatment (28). The active ingredient formononetin can exert its anti-tumor properties by regulating MAPK signaling pathway (29). TNF is a kind of cytokine with various biological effects, which can participate in regulating cell growth, differentiation, apoptosis and inducing inflammation (30). One study showed that psoralen, a natural non-toxic compound, induces apoptosis in prostate cancer by inhibiting TNF signaling pathway (31). In addition, it has found that regulating the IL-17 signaling pathway can prevent the occurrence of esophageal cancer in mice (32). Metformin regulates the progression of multiple myeloma by inhibiting HIF-1 signaling pathway (33). Thus, we found that these signaling pathways can participate in carcinogenesis, and the anti-breast cancer effect of the active ingredients in San Huang Decoction may be related to these signaling pathways. However, this study analyzed the molecular mechanism of San Huang Decoction against breast cancer, but it still needs to be confirmed by detailed experiments.

In conclusion, our study revealed the active ingredients, target genes and related signaling pathways of San Huang Decoction against breast cancer through network pharmacology. We screened a total of 39 active ingredients from San Huang Decoction. 185 potential anti-breast cancer targets of San Huang Decoction were collected from GeneCards, OMIM, PharmGkb, TTD and DrugBank databases, and 16 key targets were screened by PPI topological network analysis. In addition, GO and KEGG enrichment analysis revealed that the molecular mechanism of San Huang Decoction against breast cancer was closely related to important biochemical processes and signaling pathways, such as nuclear receptor activation, oxidative stress, PI3K-Akt signaling pathway, MAPK signaling pathway and TNF signaling pathway. The current study provides a research basis for in-depth study of San Huang Decoction in the treatment of breast cancer.

## Notes

### Competing Interest Statement

The authors have declared no competing interest.

## References

1. Fan L, Strasser-Weippl K, Li J-J, St Louis J, Finkelstein DM, Yu K-D, et al. Breast cancer in China. Lancet Oncol. 2014;15(7):e279–e89.

2. Yu L-Y, Tang J, Zhang C-M, Zeng W-J, Yan H, Li M-P, et al. New Immunotherapy Strategies in Breast Cancer. Int J Environ Res Public Health. 2017;14(1).

3. Maughan KL, Lutterbie MA, Ham PS. Treatment of breast cancer. Am Fam Physician. 2010;81(11):1339–46.

4. Xu Y, Chen X, Chen X, Bian W, Yao C, Zhang X, et al. San Huang Decoction downregulates Aurora kinase A to inhibit breast cancer cell growth and enhance chemosenstivity to anti-tumor drugs. Pathol Res Pract. 2016;212(8):696–703.

5. Xu C, Wang Y, Feng J, Qin L, Xu R, Dou Y. Effect of optimal combination of Huangqi (Radix Astragali Mongolici) and Ezhu (Rhizoma Curcumae Phaeocaulis) on proliferation and apoptosis of A549 lung cancer cells. J Tradit Chin Med. 2018;38(3):351–8.

6. Zhang X, Chen Y, Zhang T, Zhang Y. Inhibitory effect of emodin on human hepatoma cell line SMMC-7721 and its mechanism. Afr Health Sci. 2015;15(1).

7. Aggarwal BB, Yuan W, Li S, Gupta SC. Curcumin-free turmeric exhibits anti-inflammatory and anticancer activities: Identification of novel components of turmeric. Mol Nutr Food Res. 2013;57(9):1529–42.

8. Xu Y, Wang C, Chen X, Li Y, Bian W, Yao C. San Huang Decoction Targets Aurora Kinase A to Inhibit Tumor Angiogenesis in Breast Cancer. Integr Cancer Ther. 2020;19:1534735420983463.

9. Zhu ZY, Xue JX, Yu LX, Bian WH, Zhang YF, Sohn KC, et al. Reducing postsurgical exudate in breast cancer patients by using San Huang decoction to ameliorate inflammatory status: a prospective clinical trial. Curr Oncol. 2018;25(6):e507–e15.

10. Luo T-T, Lu Y, Yan S-K, Xiao X, Rong X-L, Guo J. Network Pharmacology in Research of Chinese Medicine Formula: Methodology, Application and Prospective. Chin J Integr Med. 2020;26(1):72–80.

11. Poornima P, Kumar JD, Zhao Q, Blunder M, Efferth T. Network pharmacology of cancer: From understanding of complex interactomes to the design of multi-target specific therapeutics from nature. Pharmacol Res. 2016;111:290–302.

12. Li S, Zhang B. Traditional Chinese medicine network pharmacology: theory, methodology and application. Chin J Nat Med. 2013;11(2):110–20.

13. Chen AY, Chen YC. A review of the dietary flavonoid, kaempferol on human health and cancer chemoprevention. Food Chem. 2013;138(4):2099–107.

14. Wang K, Liu X, Liu Q, Ho IH, Wei X, Yin T, et al. Hederagenin potentiated cisplatin- and paclitaxel-mediated cytotoxicity by impairing autophagy in lung cancer cells. Cell Death Dis. 2020;11(8):611.

15. Zhai T, Zhang X, Hei Z, Jin L, Han C, Ko AT, et al. Isorhamnetin Inhibits Human Gallbladder Cancer Cell Proliferation and Metastasis via PI3K/AKT Signaling Pathway Inactivation. Front Pharmacol. 2021;12:628621.

16. Xin M, Wang Y, Ren Q, Guo Y. Formononetin and metformin act synergistically to inhibit growth of MCF-7 breast cancer cells in vitro. Biomed Pharmacother. 2019;109:2084–9.

17. Chen J, Zhang X, Wang Y, Ye Y, Huang Z. Differential ability of formononetin to stimulate proliferation of endothelial cells and breast cancer cells via a feedback loop involving MicroRNA-375, RASD1, and ERα. Mol Carcinog. 2018;57(7):817–30.

18. Li T, Zhang S, Chen F, Hu J, Yuan S, Li C, et al. Formononetin ameliorates the drug resistance of Taxol resistant triple negative breast cancer by inhibiting autophagy. Am J Transl Res. 2021;13(2):497–514.

19. Olivier M, Hollstein M, Hainaut P. TP53 mutations in human cancers: origins, consequences, and clinical use. Cold Spring Harb Perspect Biol. 2010;2(1):a001008.

20. Leroy B, Anderson M, Soussi T. TP53 mutations in human cancer: database reassessment and prospects for the next decade. Hum Mutat. 2014;35(6):672–88.

21. Shahbandi A, Nguyen HD, Jackson JG. TP53 Mutations and Outcomes in Breast Cancer: Reading beyond the Headlines. Trends Cancer. 2020;6(2).

22. Duffy MJ, O’Grady S, Tang M, Crown J. MYC as a target for cancer treatment. Cancer Treat Rev. 2021;94:102154.

23. Tu R, Kang W, Yang M, Wang L, Bao Q, Chen Z, et al. USP29 coordinates MYC and HIF1α stabilization to promote tumor metabolism and progression. Oncogene. 2021;40(46):6417–29.

24. Dustin D, Gu G, Fuqua SAW. ESR1 mutations in breast cancer. Cancer. 2019;125(21):3714–28.

25. Einarsdóttir K, Darabi H, Czene K, Li Y, Low YL, Li YQ, et al. Common genetic variability in ESR1 and EGF in relation to endometrial cancer risk and survival. Br J Cancer. 2009;100(8):1358–64.

26. Noorolyai S, Shajari N, Baghbani E, Sadreddini S, Baradaran B. The relation between PI3K/AKT signalling pathway and cancer. Gene. 2019;698:120–8.

27. Kashafi E, Moradzadeh M, Mohamadkhani A, Erfanian S. Kaempferol increases apoptosis in human cervical cancer HeLa cells via PI3K/AKT and telomerase pathways. Biomed Pharmacother. 2017;89:573–7.

28. Burotto M, Chiou VL, Lee J-M, Kohn EC. The MAPK pathway across different malignancies: a new perspective. Cancer. 2014;120(22):3446–56.

29. Tay K-C, Tan LT-H, Chan CK, Hong SL, Chan K-G, Yap WH, et al. Formononetin: A Review of Its Anticancer Potentials and Mechanisms. Front Pharmacol. 2019;10:820.

30. van Horssen R, Ten Hagen TLM, Eggermont AMM. TNF-alpha in cancer treatment: molecular insights, antitumor effects, and clinical utility. Oncologist. 2006;11(4):397–408.

31. Srinivasan S, Kumar R, Koduru S, Chandramouli A, Damodaran C. Inhibiting TNF-mediated signaling: a novel therapeutic paradigm for androgen independent prostate cancer. Apoptosis. 2010;15(2):153–61.

32. Jia X-B, Zhang Q, Xu L, Yao W-J, Wei L. Effect of Nakai Leaf Flavonoids on the Prevention of Esophageal Cancer in C57BL/6J Mice by Regulating the IL-17 Signaling Pathway. Onco Targets Ther. 2020;13:6987–96.

33. Kocemba-Pilarczyk KA, Trojan S, Ostrowska B, Lasota M, Dudzik P, Kusior D, et al. Influence of metformin on HIF-1 pathway in multiple myeloma. Pharmacol Rep. 2020;72(5):1407–17.

